# Occurrence, risk assessment of antibiotics and antimicrobial resistance in *Escherichia coli* in typical rivers of Sichuan

**DOI:** 10.1101/2024.06.13.598958

**Authors:** Jingzhou Sha, Minghao Wu, Yaliang Zhou, Tao Chen, Haisha Liu, Jingjing Zhang, Wan Luo, Yi Huang, Yinshan Liu, Baoming Wang, Tao Song, Jiafu Lin

**Author notes:** Corresponding author: Tao Song and Jiafu Lin, and. Equal contribution to this paper (Jingzhou Sha, Minghao Wu, Yaliang Zhou).

## Abstract

Worldwide interest has been generated by the presence and distribution of antibiotics and antibiotic resistance genes (ARGs) in rivers. However, there was a dearth of research on the contamination of Sichuan province’s typical rivers with antibiotics. In this study, the residual level of antibiotics in 42 national/provincial sites of 9 rivers was analyzed by UPLC-MS/MS, the ecological risk level was evaluated using risk assessment method, and the drug resistance of *E. coli* in water was evaluated by Kirby-Bauer method. Redundancy analysis demonstrated how residual antibiotics affect the structure of the microbial community in the Minjiang River basin (RDA). Nine rivers all contained antibiotics. Among them, the MinJiang, TuoJiang, and JiaLingJiang rivers were severely contaminated with antibiotic concentrations ranging from 0.29 to 2233.71 ng/L. The Sichuan Basin’s antibiotic pollution level was significantly higher when compared to other Sichuan zones, which was consistent with the region’s high population density. Additionally, it was discovered that 9.77% of the *E. coli* isolated from 9 rivers had antibiotic resistance, and more than 5.8% of them had multidrug resistance. Moreover, Norfloxacin, amoxicillin, ampicillin, and tetracycline were the main risk factors for high ecological risk in 26 of the 42 monitoring sites. Additionally, there is a strong correlation between the microbial community change and residual antibiotic. These results offered some reference information regarding the distribution of antibiotics and ARGs in typical rivers in the Chinese province of Sichuan, and this study showed that more attention needs to be paid to antibiotic pollution in Sichuan’s typical rivers.

## Introduction

Antibiotics have been widely used in livestock and aquaculture for the treatment of late infections, prevention of disease, and stimulation of growth^(1, 2)^. The global antibiotics consumption in livestock and aquaculture was 63,151 tons in 2010, which was predicted to rise by 67% by 2030^(3)^. In USA, 80% of antimicrobial annual consumption was from livestock feed^(4)^. In addition, a significant amount of antibiotics are used to maintain animal productivity in low- and middle-income countries, particularly in in Brazil^(5)^, Russia^(6)^, India^(7)^ and China^(8)^. This is due to the growing demand for animal protein. However, the persistent misuse of antibiotics has led to 50–80 percent of antibiotics ending up in the environment due to their indegradability^(9, 10)^.

Multiple antibiotics, including but not limited to oxytetracycline, chlortetracycline, tetracycline, and amoxicillin, have been discovered in the environment throughout the world^(11)^. The gradual but consistent increase of antibiotics in the environment may facilitate the binding, transmission, and gene transfer of antibiotic resistance genes (ARGs), leading to drug resistance and the failure of antibiotic treatment^(12, 13)^. According to research conducted by the European Centre for Disease Prevention and Control, approximately 25,000 Europeans perish annually from drug-resistant bacterial infections^(14)^. In addition, the British government estimates that more than half a million deaths are attributable to drug-resistant bacterial infections^(15)^.

Water systems are an important reservoir for antibiotics and ARGs^(16, 17)^. The accumulation of antibiotics and ARGs in the water could act as a long-term selective pressure, thereby reducing the diversity of microorganisms and reshaping the microbial community, and further polluting the water environment ^(18, 19)^. As a result, the bioaccumulation of antibiotics in poultry, aquatic products, and vegetables will therefore directly or indirectly enter the human body via the food chain. This presented a possible threat to the ecological environment and human health. For instance, eleven antibiotics with a total concentration of 229 ng/L were detected in the drinking water source from the Yangtze River in China^(20)^. In addition, 15 types of antibiotics, including tetracyclines, quinolones, and others, were discovered in the Yellow River of China’s surface water^(21)^.

Sichuan is a populous province in China where the medical, agricultural, and aquaculture industries are developed, and the use of antibiotics is widespread. However, research on the concentration of antibiotic residue in river water, risk assessment, and its effect on water microorganisms in Sichuan Province is limited. Yang et al. and Tuo et al. found ARGs in the Funan River in Sichuan. However, only one river was investigated, which could not represent the overall antibiotics pollution in Rivers of Sichuan^(22, 23)^.

This is the first study to investigate systematically the concentration and ecological risk of antibiotic residues in typical water systems in Sichuan (9 Rivers). In addition, the level of *E. coli* resistance in water and the effects of antibiotic residues on the structure of microbial communities in bodies of water were studied in greater depth. These results could serve as a basis for assessing antibiotics pollution in Sichuan Rivers and aid policymakers in formulating the most effective anti-pollution measures.

## Materials and Methods

### Study Sampling-sites and Sample collection

In total 1-13 sample collection sites were set on nine main typical river trunk streams (Minjiang River, FuJiang River, TuoJiang River, JiaLingJiang River, QuJiang River, DaDuHe River, YaLongJiang River, JinShaJiang River, HuangHe River) in Sichuan based on the Distribution Map of Surface Water Environmental Quality Monitoring Points in Sichuan Province during the 13th Five-Year Plan Period, as well as the basin length, basin area, basin population distribution in Sichuan Province (Table 1). The collection sites were spread across 17 prefecture-level cities and 2 autonomous prefectures. While the samples were being collected, the population distribution, industrial layout, and hydrological conditions around the sampling points were also counted (Table 2).

### Sample collection and processing

The locations of water sampling sites are the left bank, the middle bank, and the right bank of the river section. The volume of water sample in each sampling point were collected for 5 L, then the collected water samples were sent to the laboratory for processing as soon as possible. 500 mL of water samples from each sampling point were mixed and 3 mixed samples were processed in parallel. Additionally, 50 mL mixed sample was collected in a sterile centrifuge tube (602051 NEST, Shanghai China), sealed, and refrigerated in an ice bag for microbial culture and screening.

### Chemical reagents/solvents and target antibiotics

Total 15 kinds of antibiotics in 6 categories were analysed in this study. Including β-lactam (β-lactams): amoxicillin (AMP), ampicillin (AMX), cephalexin (CEX), cefotaxime (CTX); Quinolones (FQs): enrofloxacin (ENR), levofloxacin (LEV), norfloxacin (NOR), moxifloxacin (MOX); Sulphonamides (SAs): sulfadiazine (SD), Sulfamethoxazole (SMZ); Tetracycline (TCs): oxytetracycline (OTC), tetracycline (TE), chlortetracycline (CTC); Amide alcohols (CRPs): chloramphenicol (CAP); Lincomide (LINs): clindamycin (CLI). Antibiotics are purchased from the China Institute for Food and Drug Control (China Institute for Food and Drug Control, China). The solvents and chemical reagents used were analytically pure or chromatographically pure. The internal standards included C13-caffeine, amoxicillin-D4, ciprofloxacin-D8, sulfamethoxazole-D4, thiabendazole-D4 and chloramphenicol-D5 were purchased from China Institute for Food and Drug Control.

### Antibiotics analysis

Pre-treatment: 500 mL water was filtered drainage fibre glass filter membrane (20191209001,NEWSTAR China). 0.5 g Na_2_EDTA and 100 ng mixed internal standard were added as recovery indicator, then pH was adjusted to 3.0 by 4M sulfuric acid. Solid phase extraction (SPE) was used to enrich antibiotics: the water sample was loaded to the activated HLB column with 6-8 mL/min extraction flow rate. Then, the container was washed with 25 mL 5% methanol water (V_methanol_: V_water_) after sample load, and the washing solution was added to the HLB column (500 mg, 6 mL). Finally, the HLB column was washed with 10 mL ultra-pure water to remove Na_2_EDTA and other residual impurities. HLB column was dried by vacuum for 120 min, and the target substance on HLB column was eluted with 8mL methanol. Finally, the volume of the target substance reached to 1ml by adding 60% methanol water (V methanol: v water). The sample solution was stored in a brown bottle and stored at −20 ℃ after filtering the sample solution with 0.22 μm filter membrane (20191209001, NEWSTAR, China), and measured on UPLC-MS/MS (UPLC:A-30,Japan; MS/MS:ABSciex Qtrap3200, Agilent, USA).

The optimization of mass spectrometry conditions includes ion source mode, cone hole voltage and parent ion, collision energy and sub-ion. Chromatographic column: AthenaC18-WP (2.1×150 mm, 3 μm); injection volume: 5 μL; mobile phase flow rate: 0.5 mL / min; column temperature: 40 ℃. Mobile phase A is 0.1% formic acid, mobile phase B is acetonitrile. Chloramphenicol is MRM negative ion mode and other 14 antibiotics are MRM positive ion mode. Mass spectrometry conditions: UPLC-MS/MS adopts multiple reaction mode, ESI ion source, drying gas temperature and flow rate are 325 ℃ and 6ml/min, respectively. Sheath gas flow rate is 11L / min, ionization voltage is 2.5 kV, atomizer pressure is 45 psi.

### Environmental risk assessment

The ecological risk quotient (RQ_Ecotox_) of antibiotic residues in the environment was evaluated by risk quotient method based on EU risk assessment technical guidance. The formula is as follows: RQ_Ecotox_=MEC/PNEC_Ecotox_ (Note: It is assumed that there is no “synergism” or “antagonism” between antibiotics in risk assessment). MEC is the measured environmental concentrations of pollutants in the environment. PNEC_Ecotox_ is the predicted no-effect concentration, which is the toxicological data (Table 3) of the most sensitive species to target antibiotics. RQ_S_ < 0.1 is low risk grade, RQ_S_ < 1 is medium risk grade, RQ_S_ ≥ 1 is high risk grade.

### Screening/identification of Enterobacter and analysis of bacterial drug resistance

*E. coli* was isolated and screened based on the method reported by Ibrahim et al(24). Furthermore, identification and screening of *E. coli* were carried out through McConkey medium culture-Gram stain observation - fermentation experiment - oxidase experiment - indole experiment, and the *E. coli* conforming to the identification results were stored for later use. 0.1 mL of river water was evenly coated on MacConkey AGAR medium for culture. Typical *E. coli* colonies (brick red) were selected for re-purification. The purified bacteria were preserved, and the next biochemical test (fermentation test, oxidase test and indole test) was carried out. The strain with positive fermentation test, negative oxidase test and positive indole test was *Escherichia coli*.

The sensitivity of *E. coli* to four antibiotics was analysed by AGAR diffusion method (K-B method), namely gentamicin (10 μg), tetracycline (30 μg), levofloxacin (5 μg) and ofloxacin (5 μg). Susceptibilities of enterobacteria to antibiotics can be divided into susceptible (S), Intermediate (I) and Resistant (R). Sensitivity is determined according to the CLSI implementation standard (2019 version)

## Results

### Analysis of residual characteristics of antibiotics in typical rivers of Sichuan

This study included 9 typical rivers in Sichuan (Fig. S1), which were MinJiang River, DaDuHe River, JinSha River, TuoJiang River, JiaLingJiang River, QuJiang River, YalongJiang River, FuJiang River and HuangHe River. The colorful dots in the map indicated monitoring sites (Fig. S1). All 15 antibiotics were successfully detected, and the detection rate was from 2.38% (CTX) to 83.33% (LEV) (Table 4). The concentration of antibiotics in rivers varied. The top three polluted rivers were MinJiang (938 ng/L), TuoJiang (177 ng/L), and JiaLingJiang (127 ng/L) (Fig. 1). Moreover, the antibiotics concentration varied dramatically between monitoring sites in the same river (Fig. 1). The concentration of antibiotic pollution increased gradually from the upper to lower reaches of the river (Fig. S2). Specifically, in the Jinsha River, Minjiang River, Dadu River, Tuojiang River, Qujiang River, Yalong River, and Fujiang River, where the cumulative concentration of antibiotics differed significantly between the upper and middle reaches (Fig. S2, P<0.05). Based on the distribution map of antibiotic concentrations (Fig. S2), the monitoring sites with the most pollution were all concentrated in the Sichuan Basin, and the findings were in line with the local rivers’ typical antibiotic use patterns, industrial distribution, and population density.

**Figure 1.**
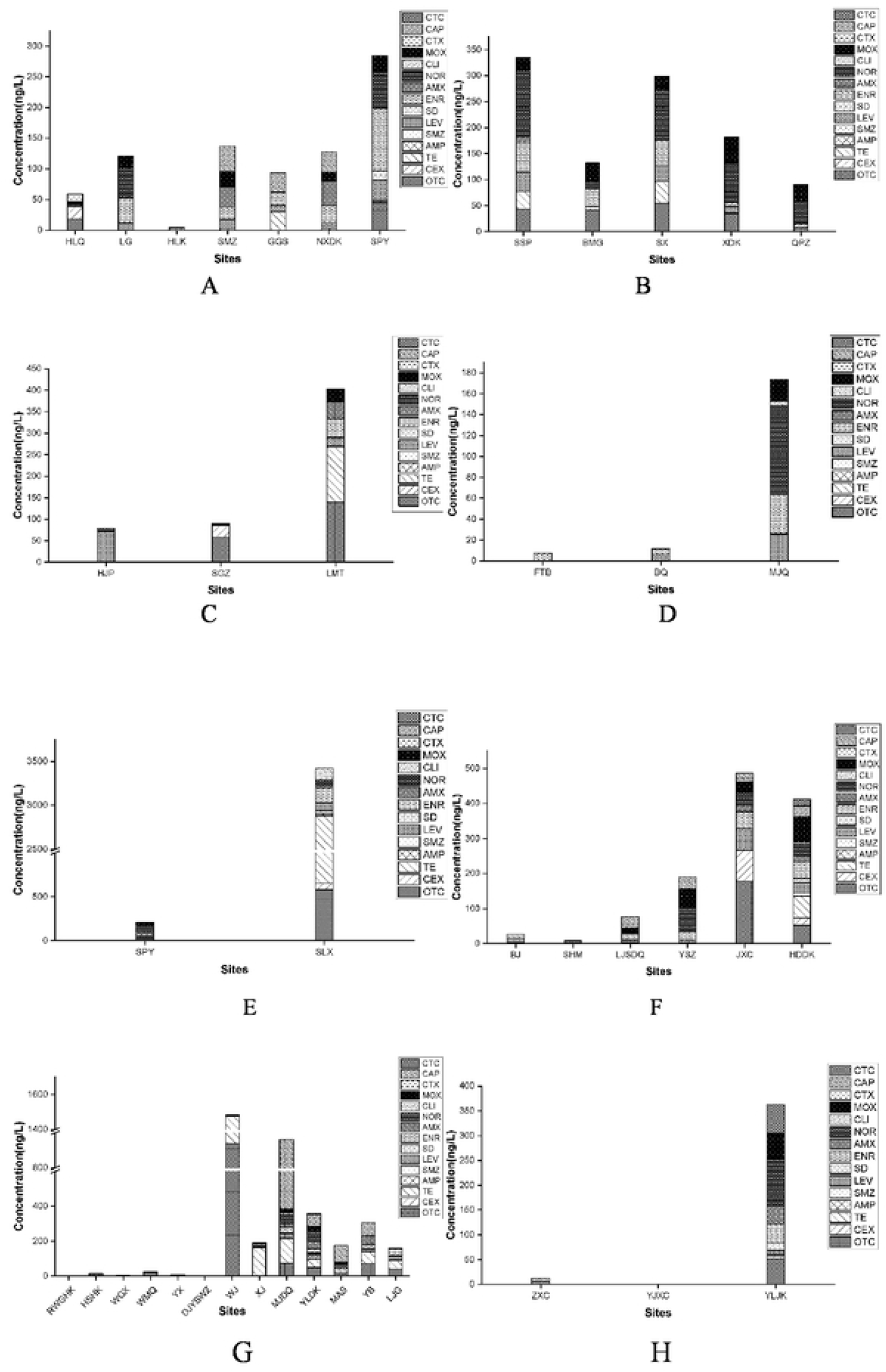
Antibiotics concentration at monitoring sitesin 9 Rivers. A, JinShaJiang River; B, Jialingjiang River; C, Daduhe River; D, Fujiang River; E, Qujiang River; F, Tuojiang River; G, Minjiang River; H, Yalongjiang River-Huanghe River. TCs, Tetracycline; OTC, oxytetracycline; CTC, chlortetracycline; TE, tetracycline; AMP, ampicillin; AMX, amoxicillin; CEX, cephalexin; CTX, cefotaxime; SAs, Sulfonamides; SD, sulfadiazine; SMZ, Sulfamethoxazole; FQs, Quinolones; ENR, enrofloxacin; LEV, levofloxacin; NOR, norfloxacin; MOX, moxifloxacin; LINs, Lincomide; CLI, clindamycin; CRPs, Amide alcohols; CAP, chloramphenicol.

### Antimicrobial resistance of *E. coli* in rivers

Antibiotic pollution is not only harmful to the ecological balance but also to people’s health. Because of the long-lasting antibiotic residue in the water, the bacteria may become resistant to these drugs, complicating medical treatment, and endangering the patient’s life. To determine the level of antibiotic resistance, 695 *E. coli* were isolated from typical rivers and then treated with 4 distinct antibiotics (CN: Gentamicin, LEV: Levofloxacin, TE: Tetracycline, and OFX: Ofloxacin). 9.77% of *E. coli* strains are resistant to at least one antibiotic and the proportion of *E. coli* resistant to antibiotics decreased because of increased antibiotic treatment (Fig. 2A). 8.63% were resistant to one antibiotic, and 0.57% were resistant to two. Multidrug-resistant bacteria (MDR) constituted only 0.43% (Fig. 2A), and only one strain (SHM-14) was resistant to four antibiotics (CN, LEV, TE and OFX). Additionally, *E. coli* exhibits varying levels of resistance to these four antibiotics (Table 5). TE was the most resistant antibiotic, as indicated by its 6.1% resistant ratio and 7.34 intermediate value (Fig. 2B). As stated previously, TE was the most prevalent antibiotic in typical rivers; this may explain why it is the antibiotic with the highest *E. coli* resistance. Furthermore, it was discovered that the YaLongJiang River and the HuangHe River had the highest and lowest levels of antibiotic resistance, respectively.

**Figure 2.**
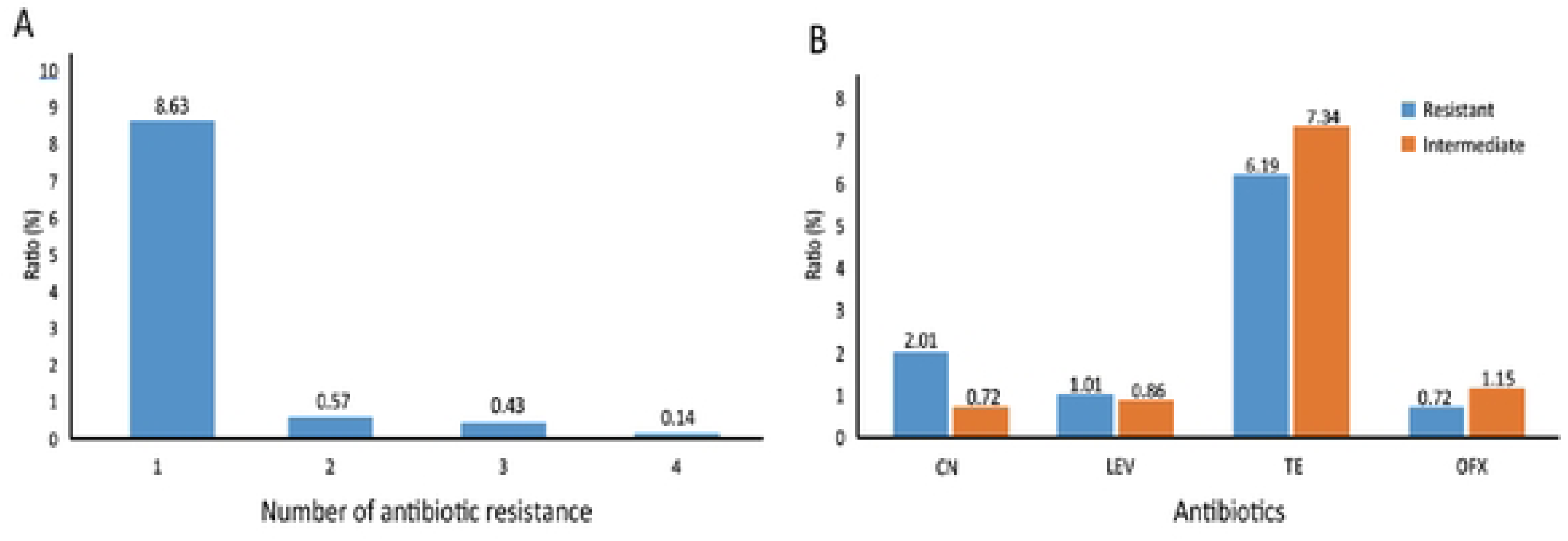
Resistance of *E. coli* to four antibiotics. A, Resistance of 695 Escherichia coli to four antibiotics. B, The proportion of different antibiotic-resistant bacteria in the population of Escherichia coli. CN, Gentamicin; LEV, Levofloxacin; TE, Tetracycline; OFX: Ofloxacin.

### Risk assessment of typical rivers

The distribution of antibiotics and drug resistant bacteria in rivers could damage the ecological balance. Therefore, ecological risk assessment of antibiotic residues in typical rivers in Sichuan was further carried out. Bacteria, algae, and water fleas were chosen as the study’s objects of analysis due to their dominance in those Rivers. As depicted in Figure 3, antibiotic residues in 26 of the 42 monitoring sites posed a high risk, while 10 posed a medium risk. The most significant environmental risk factors in the Sichuan River system were NOR, AMX, AMP, and TE. The highest level was shared by NOR and AMP (38.09%), followed by AMX (30.95%), TE (16.67%), ENR (9.52%), CAP (9.52%), and OTC (4.76%). In addition, CTC, CLI, and SMZ were at the bottom (2.38%). RQEcotox values for 15 antibiotics ranged from 0 to 72.94. CTX posed the lowest ecological risk among 42 monitoring sites, with the lowest mean RQEcotox value (0.001). In contrast, AMP and AMX were found to display the greatest ecological risk, as indicated by the highest average (4.066 and 2.433) values. Additionally, the ecological risk varied in the different typical rivers. HuangHe River and FuJiang river exhibited the lowest ecological risk (Fig. 3). In contrast, organisms in other seven rivers were heavily contaminated by antibiotics. In addition, the risk level varied significantly between upstream and downstream regions. In the middle and downstream areas, where most monitoring sites were labelled as highest risk and the ecological risk was greatest. As stated previously, the population is more concentrated in the basin’s middle and lower reaches. This indicates that the ecological risk posed by antibiotics was strongly linked to human activity.

**Figure 3.**
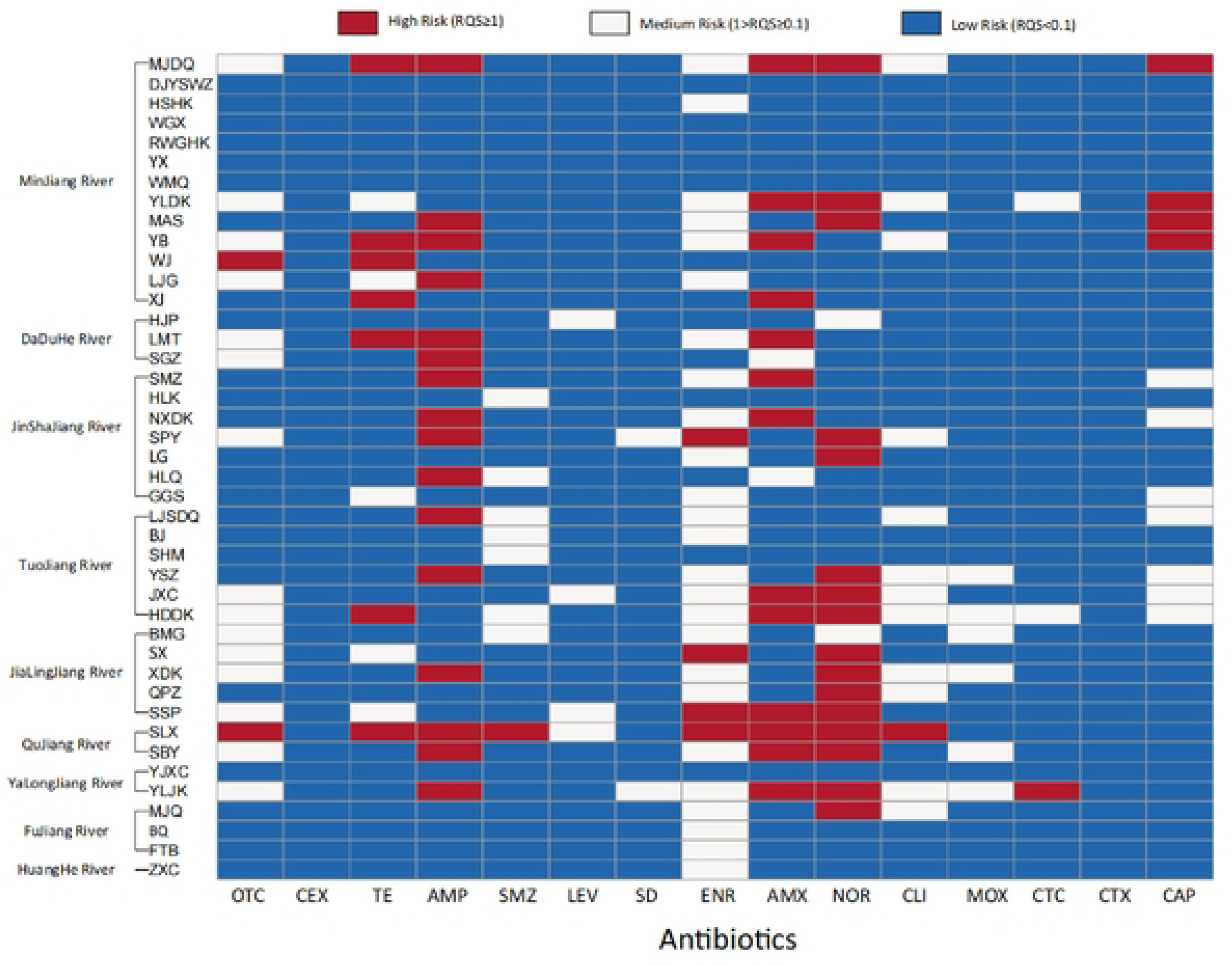
Risk assessment of antibiotic at monitoring. TCs, Tetracycline; OTC, oxytetracycline; CTC, chlortetracycline; TE, tetracycline; AMP, ampicillin; AMX, amoxicillin; CEX, cephalexin; CTX, cefotaxime; SAs, Sulfonamides; SD, sulfadiazine; SMZ, Sulfamethoxazole; FQs, Quinolones; ENR, enrofloxacin; LEV, levofloxacin; NOR, norfloxacin; MOX, moxifloxacin; LINs, Lincomide; CLI, clindamycin; CRPs, Amide alcohols; CAP, chloramphenicol.

### Microbial Diversity and Community structure of Minjiang River

To further explore the influence of antibiotics to the microbial diversity, the microorganism from 51 water samples of 13 monitoring site were sequenced. The RWGHX monitoring size had the highest Species (Fig. 4A), Shannon (Fig. 4B), Simpson (Fig. 4C), and Chao 1 (Fig. 4D) indices of microbial diversity and richness (4585, 9.75, 9.94 and 5427). This means the ecological environment in RWGHX was the best. In contrast, WGX possessed the lowest value of Species (1098), Shannon (6.66), Simpson (0.95), and Chao 1 (1332). Furthermore, the similarity of the microbial community was explored. There was 179 same operational taxonomic units (OUT) among the 13-monitoring site (Fig. 5A). Additional OUTs could be found in the site. *Proteobacteria* was found to be the dominant population in all the samples, with an average proportion range of 24%-71.25%, followed by *Actinobacteria* (1.8%-38.17%) and *Bacteroidetes* (12.04%-35.19%), respectively (Fig. 5B). Our study demonstrated that antibiotics pollution can reduce proteobacteria abundance, as the proportion of proteobacteria was greater than 50 percent in samples with a lower antibiotic concentration (DJYSWZ, YX, WGX and RWGHK).

**Figure 4.**
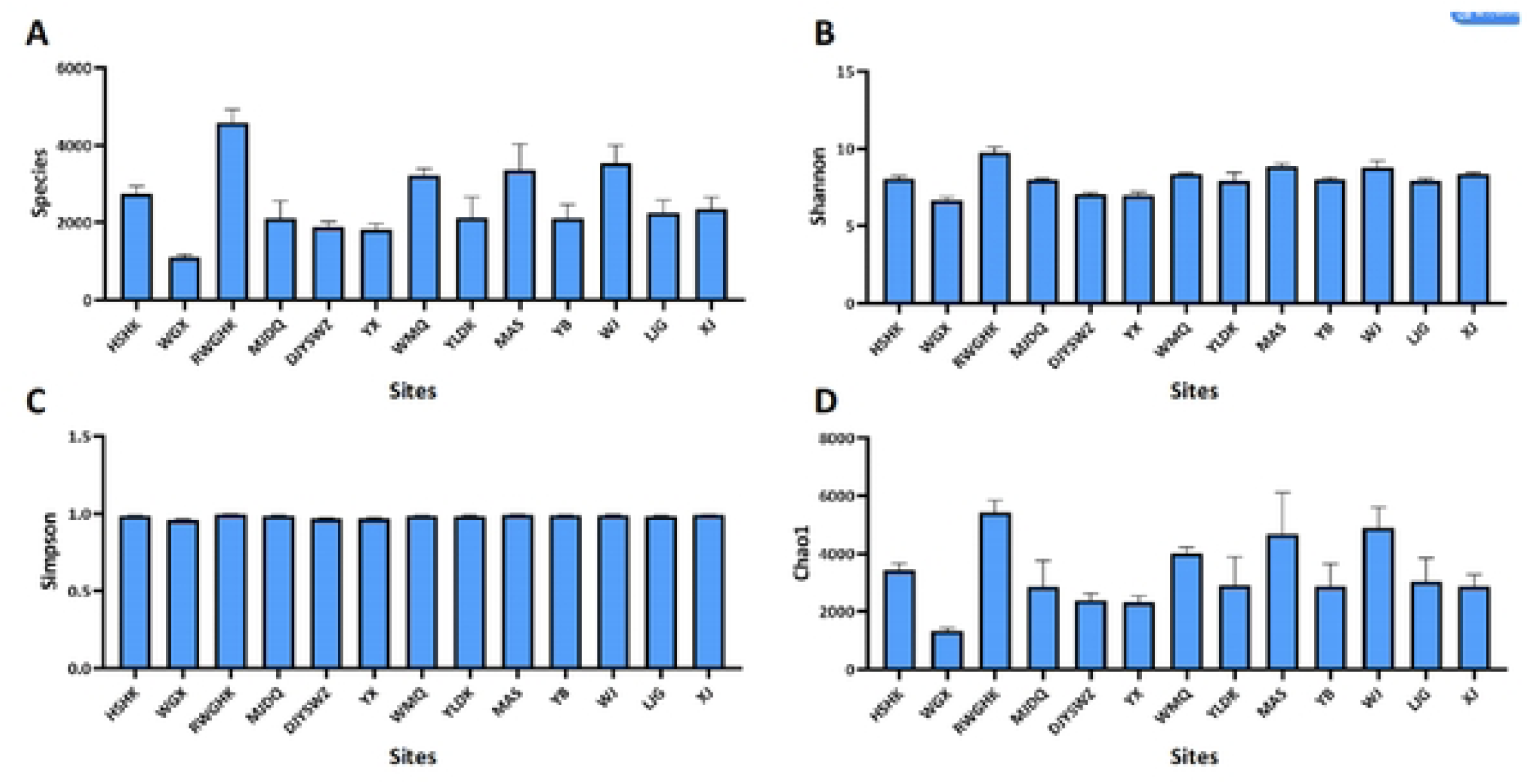
α-Diversity data of monitoring sites in Minjiang River. A, Species value. B, Shannon value. C, Simpson value. D, Chao 1 value.

**Figure 5.**
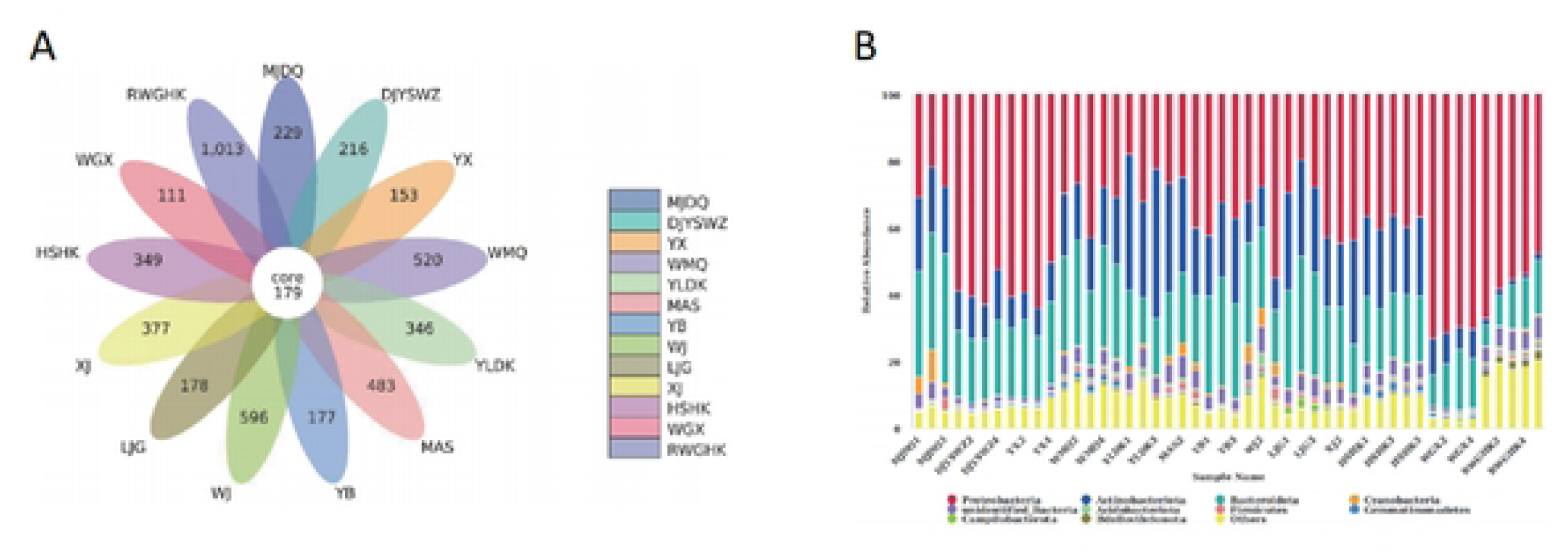
Operational taxonomic units and Taxonomic composition distribution of MinJiang River. A, Venn diagrams that showing the shared and distinct OTUs in sites of Minjiang River. B, Taxonomic composition distribution histograms in monitoring sites at phylum. OTUs, operational taxonomic units.

To explore the relationship between antibiotics and microbial community in more details, redundancy analysis (RDA) was included in the study (Fig. 6). Total 6 antibiotics (OTC, CAP, LEV, NOR, SD and CTC) were selected based on the result from Monte Carlo test. According to the RAD analysis, WMQ, DJYSWZ and RWGHK showed similar microbial community. This may attribute to closed location of those monitoring sites, which resulted in the similar carbon and nitrogen content. Additionally, LEV, SD, NOR, CTC and CAP were found to significantly affect the microbial community. The negative association was found between *Proteobacteria* and LEV[-0.756_Proteobacteria_(p<0.01)] and SD[−0.583_Proteobacteria_(p<0.05)]. On the contrary, LEV[0.744_Actinobacteriota_ (p<0.01)], SD[0.71_Actinobacteriota_ (p<0.01)], NOR[0.563_Actinobacteriota_ (p<0.05)] and CTC[0.594_Actinobacteriota_ (p<0.05)] present positive association to *Actinobacteriota.* Besides, CAP was positively associated to the *Bacteroidota* [0.555_Bacteroidota_ (p<0.01)] and *Cyanobacteria*[0.694_Cyanobacteria_ (p<0.01)]. Most antibiotics present positive association to the microbial community.

**Figure 6.**
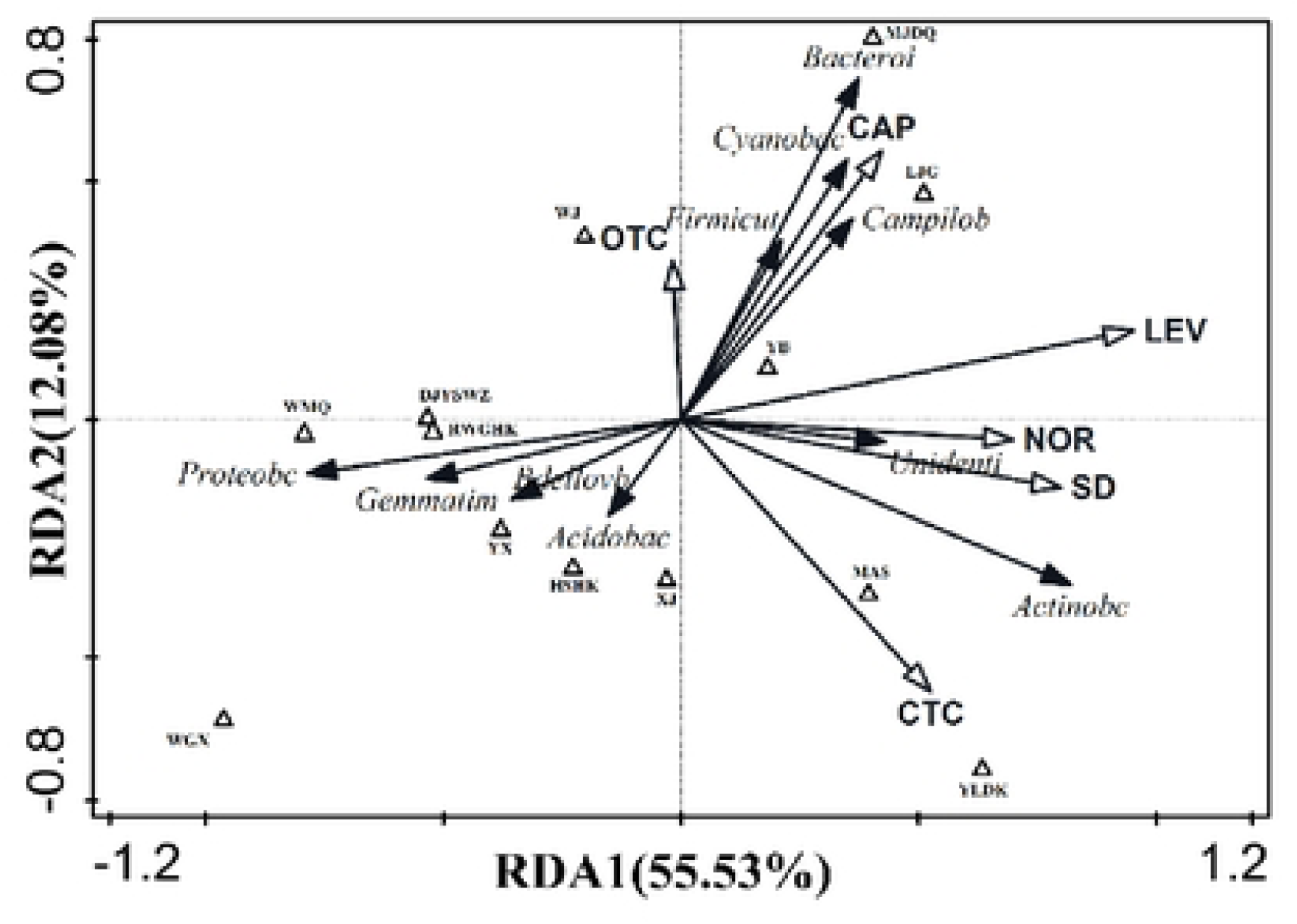
Redundancy analysis (RDA) of antibiotics residual and bacterial communities. Triangle represents each sample. Antibiotics and bacterial (phylum) were indicated by hollow arrows and Solid arrows.

## Discussion

Antibiotic pollution in rivers is a significant issue that can have negative consequences for both human health and the environment. This study investigated the level of antibiotic residues and ecological risks induced by antibiotics. Additionally, the effects of antibiotic residues on *E. coli* antibiotics resistance, as well as on the microbial community in the typical water system of Sichuan in were further assessed. Our research revealed that all 15 antibiotics were present in nine typical Sichuan rivers in concentrations ranging from 0.29 ng/L to 2233.71 ng/L and the main pollutant in the rivers was FQs. Furthermore, drug resistance was found in the 9.77% of *E. coli* and more than 5.8% drug resistant *E. coli* was MDR. Moreover, 26 of 42 monitoring sites were in the high ecological risk level and NOR, AMP, AMX and TE were identified as the prominent risk factors in 9 typical rivers of Sichuan. Finally, it was discovered that there was a strong correlation between the microbial community and LEV, SD, NOR, TCT, and CAP, suggesting that those antibiotics may have an impact on the ecological balance of the MinJiang River.

Antibiotics, such as TCs and FQs, has been commonly used in both human and animal medicine. Numerous studies have proven the existence of antibiotics in the environment, including in rivers^(25–27)^. In our study, 15 antibiotics had been included and were all found in 9 rivers of Sichuan province. The average concentration of 15 antibiotics in 9 rivers was comparable to the concentrations of 36 frequently detected antibiotics in 58 rivers as determined by another systemic analysis study^(28)^. However, the detection rate of TCs and SAs in our study was much lower (61.9% and 30%) than Haihe River ^(29)^ and Yangtze River of China^(30)^. This difference may be attributable to the low prevalence of TCs in Sichuan as a result of the obvious side effect in the clinic and the strong adsorption action of the particulate matter and sediment in the water^(31)^.

As mentioned above, the antibiotics included in the study were commonly used in human and animals. Therefore, the correlation between antibiotics concentration and population size of human or animals are essential. In our study, the same antibiotic concentration was found to be varied between rivers. The population size should be responsible for the antibiotic’s disparity between rivers. For example, the antibiotics concentration in Minjiang River was much higher than HuangHe river. Interestingly, along the Minjiang River, the permanent human population was 25 million^(32)^, whereas along the HuangHe River, it was only 0.4 million^(33)^. More people use antibiotics, which could lead to a high concentration of antibiotic residues in rivers. Furthermore, the antibiotic concentration within the same river was different between monitoring sites. The antibiotic concentration disparity were also found in China’s YongJiang River^(34)^ and FenHe River^(35)^. Those studies also demonstrated the higher human population size along the downstream of the river could increase the antibiotic concentration. Another factor could affect the antibiotic usage is animal usage. The animal husbandry is well developed in Sichuan basin because of special geographical environment. Nearly 85% of the people living in the Sichuan province reside in the Sichuan Basin, which has led to an enormous amount of farming. But the intensive farming rate was less than 50%. This indicates that more than 50% of farming activity was carried out by small-scale or individual farmers, which led to an overuse of antibiotic feed due to the high regulatory difficulty. Furthermore, it is challenging to properly dispose of agricultural waste, which enters the environment through direct discharge and causes residual antibiotics to end up in the water. These findings suggested that the most direct and efficient way to further reduce antibiotic contamination will be to lower the concentration of antibiotics in wastewater from livestock and aquaculture.

Long-term antibiotic pollution could induce drug resistance, which could be detrimental to human beings. The MDR phenomenon could be found in our study, however, the MDR rate was lower than other rivers in China, such as TongJiang River^(36)^ (MDRR=87.5%) and JiuLongJiang River^(37)^ (MDRR=70.6%). The disparate result could attribute to the antibiotic types. Those studies included highest prevalent antibiotics in the analysis, however, some antibiotics in our study was relatively less popularized. In addition, the Chinese government enacted more stringent environmental protection policies in recent time, including the prohibition of antibiotics as feed additives and the improvement of sewage discharge standards. These factors may explain the low prevalence of MDR in our study. Additionally, our study proved that antibiotics pollution in rivers could damage the ecological balance and AMP and AMX were found to display the greatest ecological risk. Previous research demonstrated that AMP and AMX may be toxic to microcystic aeruginosa at extremely low concentrations (3.7 ng/L and 0.31 ng/L)^(38, 39)^. Despite the fact that the half-life of lactam antibiotics in water environment was only 1-2 hours, these data demonstrated that the ecological risk posed by these antibiotics warrants sufficient consideration^(40)^. Furthermore, the ecological risk induced by antibiotics pollution was found to be associated to the changes in microbial community structure and most antibiotics displayed positive association to the microbial community in our study. The similar findings were also found in other studies. For example, LEV in low concentration still could inhibit *Azospirillum* and impair the ability of microorganisms to fix nitrogen^(41)^. Similarly, Vanesa et al. reported that CTC was toxic to the microbial community in soil of France^(42)^. All in all, those results proved that residual antibiotics can be used as environmental selective pressure to change microbial community structure and affect microbial composition.

## Conclusion

Our research explored the potential mechanism of antibiotic contamination in rivers and emphasized the hazard to human health. Those results pave the theoretical foundation on how to stop the release of antibiotic pollutants and prevent the spread of drug-resistant *E. coli*, and lessen the environmental harm caused by antibiotic residues.

## Acknowledgement

This research was supported by Sichuan Ecological Environment Protection Science and Technology Project (No.2019HB14).

